# Patterns of *Mustelid gammaherpesvirus 1* (MusGHV-1) genital reactivation linked to stressors in adult European badgers (*Meles meles*)

**DOI:** 10.1101/2021.01.19.427370

**Authors:** Ming-shan Tsai, Sarah Francois, Chris Newman, David W. Macdonald, Christina D. Buesching

## Abstract

Herpesvirus infections are common and mostly asymptomatic in vertebrates, but can result in impaired reproduction. It is therefore important to understand infection patterns and associated risk factors, particularly the effects of different stressors. Here we use Mustelid gammaherpesvirus 1 (MusGHV-1) infection in European badgers (*Meles meles*) as a host-pathogen wildlife model to study the effects of a variety of demographic, social, physiological and environmental stressors on viral reactivation in the genital tract. We collected 251 genital swabs from 151 free-ranging individuals across 3 trapping seasons (spring, summer and autumn). We screened for MusGHV-1 using PCR and explored possible links between genital MusGHV-1 reactivation and stressors, and their interactions, using logistic regression. In adults, reactivation was more likely in males, especially those in poorer body condition during summer. In females, reactivation was more likely when living in social groups comprising a higher percentage of cubs, but counter to our predictions, recent lactation appeared not influential. In relation to age, reactivation was more common in individuals over 8 years old than among prime age adults, and among juveniles (<2 years old), especially females and individuals in better body condition, likely due to early puberty. Environmentally, reactivation was more prevalent in summer when food abundance is typically low. Our results evidence age effects on MusGHV-1 reactivation; in juveniles MusGHV-1 shedding in the genital tract is likely related to primary infection, while in adults, genital MusGHV-1 reactivation from latency was associated with aging, social and/ or environmental stress.

**Importance:** The immuno-suppressive effects of elevated stress levels facilitate disease development, and can ultimately cause host extinction at the population level, especially where diseases are transmitted sexually. The impacts of stress on host-pathogen dynamics through disease, however, are still poorly understood outside the laboratory or captive environments. Our study provides rare evidence from a free-ranging wild mammal population that the infection dynamics of a common and sexually transmittable gammaherpesvirus are linked to demographic, social, physiological and environmental stress. We propose that the effects of stressors on STIs and viral reactivation are an important factor to be taken into account in conservation efforts when working with vulnerable wildlife populations.

## Introduction

Herpesvirus infection is common in vertebrates with most vertebrate herpesviruses belonging to 4 subfamilies, the *Alphaherpesvirinae, Betaherpesvirinae*, *Gammaherpesvirinae* and *Deltaherpesvirinae* (1, 2). Herpesvirus species are generally host-specific, but cross-species transmission is more frequent than previously assumed (3, 4). After primary infection, the herpesvirus enters a latent stage in the host cell (e.g. lymphocytes in gammaherpesvirus infection), and can be reactivated repeatedly throughout life by stress (5, 6), trauma (e.g. surgery:(7)) or primary co-infection with other pathogens (8). Reactivation is a process of viral lytic infection, which involves virus replication within the host cell, eventually destroying the cell and releasing infectious virions. Reactivation of herpesviruses usually occurs in the epithelial cells of mucosa that function as portals for external contact (e.g., mouth, nose, eyes and genital tract), thus facilitating transmission. Reactivation is, however, typically asymptomatic or induces only mild disease, but can also promote development of severe diseases like cancer (9), depending on strain pathogenicity (10) and co-infection with other pathogens causing immunodeficiency (e.g. Human herpesvirus 8 and HIV (11)), and is associated with a higher risk of contracting co-infection with additional pathogens with high virulence (e.g. *Chlamydia pecorum* infection in koalas suffering from gammaherpesvirus reactivation (12)).

Chronic stress has proven a significant risk factor, causing immune system dysregulation. where corticosteroids inhibit the pro-inflammatory cytokine responses, allowing the virus to (re-)activate and undergo lytic proliferation unchecked (13). This link between elevated corticosteroid levels and herpesvirus reactivation has been proven experimentally (horses: (14); captive reindeer: (15) and through observation (e.g. humans: (6, 16); captive Grévy’s zebras (*Equus grevyi*): (5), but has not been investigated in free-living wildlife populations.

Here, we use European badgers (*Meles meles*) as a wildlife model to investigate how different risk factors and stressors affect herpesvirus reactivation. Badgers are seasonally breeding mustelids that are commonly infected with the *Mustelid gammaherpesvirus 1* (MusGHV-1: a large double-stranded DNA virus belonging to the *Gammaherpevirinae*, genus *Percavirus*), where prevalence of viral DNA in blood samples can reach up to 100% in the UK and in Ireland (17, 18), and 55% - 82.5% in genital swab samples (19, 20). Gammaherpesvirus reactivation can cause severe disease in humans (21, 22) and domestic animals (23–25), and has increasingly been associated with illness in wildlife species (26–29). In badgers, previous research has linked otherwise asymptomatic MusGHV-1 reactivation in genital tracts to impaired female reproductive performance (20) and indicates that during the main mating season, adult males are at particular risk of genital MusGHV-1 reactivation. Nevertheless, the impact of social, physiological and environmental stress on MusGHV-1 reactivation has thus far not been investigated. Badgers are subject to a variety of stressors: faecal corticoid levels indicate that badgers experience seasonal variation in stress levels (30, 31), which may be due to variation in food availability (30, 31) and can result in mortality (32, 33). Sociologically, higher social group density is associated with female reproductive suppression (34, 35), reduced body condition and fecundity (36), and increased bite wounding among male badgers (37). Furthermore, aging reduces tolerance to stress (38), specifically altering the balance of innate and acquired immunity in badgers (39), and increasing their risk of herpesvirus reactivation (20), as also observed in other carnivora species (40, 41), sometimes resulting in chronic and continuous herpesvirus reactivation (42). Therefore metrics of body condition, especially reduced body-condition as a consequence of recent lactation (30, 43), can indicate that the individual may be experiencing physiological stress (30).

To evaluate the impact of potential stressors on genital MusGHV-1 reactivation, we conducted population-wide molecular screening using genital swabs taken from a free-ranging badger population in the south of England across 3 seasons (spring, summer and autumn). We investigated whether environmental factors (i.e., season and social group size), host demographic parameters (i.e., sex, age and lactation), and host health (i.e., body condition) affect risk of genital MusGHV-1 reactivation.

## Materials and methods

### Field data and sample collection

Samples were collected from 151 individual live-trapped badgers in Wytham Woods, Oxfordshire, UK (51°46’26”N, 1°19’19”W; caught in May, September and November 2018 following the methodology described in Macdonald et al. (44); for details see Table 1). All trapping and animal sampling protocols were approved by the University of Oxford’ Animal Welfare and Ethical Review Board. Trapping was conducted under Natural England license (currently 2019-38863, Badger Act 1992) and all animal handling procedures were carried out by qualified Personal Individual License (PIL) holders under Home Office license (current PPL 30/3379, Animals (Scientific Procedures) Act 1986). For each capture, we recorded sex, sett (i.e., communal den used by a badger social group) of capture, body condition score (BCS, categorized as 1= very thin to 5= very fat), and lactational status (determined by teat measurements of females in spring: (45)). Because each badger in Wytham is given an individual tattoo at first capture (usually as a cub (46)), exact age (in years) was known for most (243 of 251) animals in the dataset. For the remaining 8 badgers first caught as adults, age was inferred by toothwear according to the method described Bright Ross et al. (47). We defined 4 age classes: i) juveniles < 2 years old (cubs and yearlings were combined to increase sample sizes, as there was no difference between Mus GHV-1 prevalence in cubs and yearlings: Fisher’s exact test: p-value=0.7449); and - based on sex-steroid levels (48); ii) young adults: 2≤ x <5 years old; iii) old adults: 5≤ x <8 years old; iv) very old adults: ≥ 8 years old. The number of cubs and adults resident in each sett was estimated using minimum number alive (MNA) estimates (44, 47).

**Table 1:**
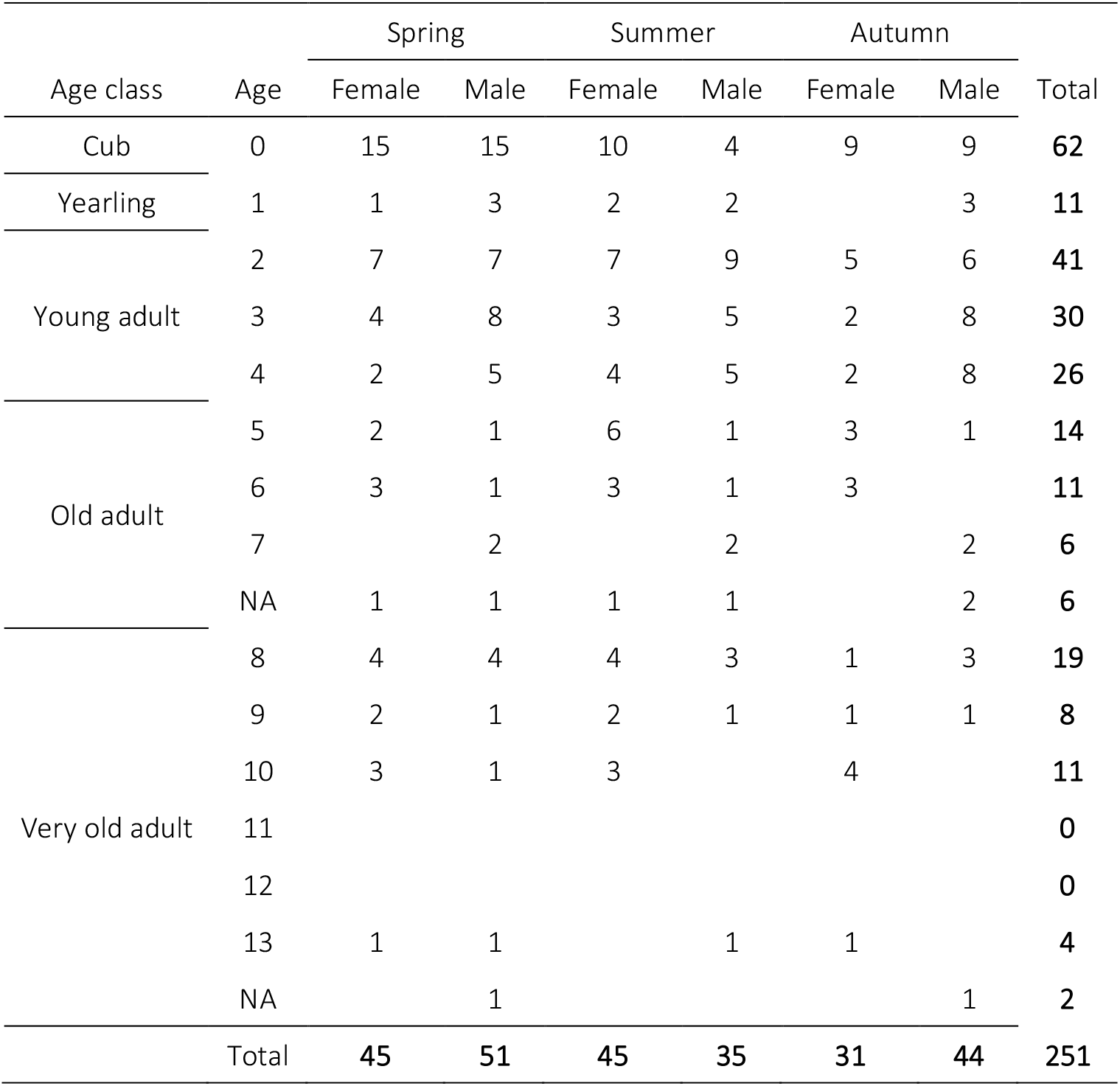
Details of swab sampling

Sterile cotton tops with wooden shafts were used to swab the genital tracts of all females (cubs and adults) and all males (except for very small male cubs in spring for animal welfare reasons), and stored in 2 ml sterile microcentrifuge tubes. All samples were frozen and stored at −20°C immediately after sampling. Badgers were released at their site of capture on the same day, after full recovery from aneasthesia.

### DNA extraction and purification

Each swab was reconstituted with 400μl sterile double distilled water and vortexed gently at room temperature for 10 minutes. A 200μl aliquot was taken from the reconstituted swab fluids, and viral DNA was extracted and purified using a commercial kit (DNeasy Blood and Tissue Kit, Qiagen) following manufacturer’s instructions. Purified DNA was then eluted in 100μl of the provided buffer.

### Screening for MusGHV-1 DNA using polymerase chain reaction (PCR) and sequencing

The purified DNA was screened using a MusGHV-1-specific primer pair designed by King et al. 2004 (18), targeting 281 base pairs of the partial DNApol gene. For each reaction, a total of 20 μl PCR solution was mixed with 10μl HotStartTaq Master Mix (Qiagen, containing 1 unit of HotStartTaq DNA Polymerase, 12μM of MgCl_2_ and 1.6μM of each dNTP), 0.5 μM of each primer, and 2μl CoralLoad gel loading dye and 5 μl DNA template. Amplification conditions were kept at 95°C for 5 mins to activate DNA polymerase, followed by 45 cycles of denaturation at 95°C for 45 seconds, primer annealing at 60°C for 45 seconds, and chain elongation at 72°C for 1 minute, followed by a final extension at 72°C for 10 minutes. Finally, the PCR products were loaded in 2% agarose gel to check the amplification results under UV light. Samples with positive results were then amplified again with substituted front primer (5’ CCA AGC AGT GCA TAG GAG GT 3’) to generate longer sequences (771 base pairs). PCR products were then purified and sent for genotyping using Sanger sequencing to confirm the identity of produced amplicon. Sequences returned were then aligned by Clustal W method (49) and analyzed for variation using MEGA X (10.1.7) (50). Representative sequences were selected and published on GenBank under accession number MT332100 and MT332101 assigned.

### Statistical analysis

Statistical analyses were performed with the R and R Studio software (version 1.21335) (51). Prevalence of genital MusGHV-1 DNA was calculated by dividing the number of PCR positive cases by the total number of tested cases, and 5% upper and lower confidence intervals were calculated using the Wilson method (52). Logistic regression (glmer function, R package lme4) with badger identity (tattoo) number as a random effect was used to measure univariate effects of MusGHV-1 reactivation in genital tracts with season, sex, age, age class, BCS, number of residents per sett, and percentage of cubs per social group (calculated by sett), where we categorized these data on percentage of cubs per group into low and high using 30% as the dividing point according to distribution of the data (Figure S1). Effects of lactation were analyzed using Fisher’s exact tests due to low sample sizes and presented using odds ratios. The final multivariable model was selected through the manual backwards selection method. Model residual diagnostics were conducted using R package DHARMa (version 0.3.3.0). Model fit was established using area-under-receiver-operating characteristics (AUC) (53). Kruskal-Wallis tests were used to compare genital MusGHV-1 positive and negative individuals of different BCS. Because juveniles (especially cubs) are generally thinner than adults, and thus have a different body condition distribution (Figure S2), we calculated a body condition index (BCI) as ln(body weight)/ln(body length) (36)). We analysed the association of MusGHV-1 reactivation and individual BCS for juveniles and adults separately. Linear models (lm function, R package lme4) were used to assess the association of age and MusGHV-1 reactivation prevalence.

## Results

The overall prevalence of genital MusGHV-1 reactivation was 35.9% (90/251, 95% CI: 30.2% - 42.0%), and prevalence was generally higher in summer (45%, 36/80) than in spring (34.4%, 33/96) and autumn (28%, 21/75) (Table 2; Figure 1).

**Table 2:** Overview of genital MusGHV-1 reactivation prevalence and univariate logistic regression analysis. Formula: MusGHV ~ Variate + (1|Tattoo); number of observations: 251; groups by tattoo number: 150

|  | Positive | Total | Prevalence | Prevalence 95% CI | Odds ratio (OR) | OR 95% CI | P-value |
| --- | --- | --- | --- | --- | --- | --- | --- |
| Sex |  |  |  |  |  |  |  |
| Male | 45 | 130 | 34.62% | 27% - 43.1% | 0.59 | 0.53 - 1.52 | 0.679 |
| Female | 45 | 121 | 37.19% | 29.1% - 46.1% |  |  |  |
| Season |  |  |  |  |  |  |  |
| Spring | 33 | 96 | 34.38% | 25.6% - 44.3% | 1.37 | 0.7 - 2.69 | 0.362 |
| Summer | 36 | 80 | 45.00% | 34.6% - 55.9% | **2.16** | **1.08 - 4.33** | **0.03** |
| Autumn | 21 | 75 | 28.00% | 19.1% - 39% |  |  |  |
| Age class |  |  |  |  |  |  |  |
| Juvenile (<2 years old) | 34 | 73 | 46.58% | 35.6% - 57.9% | **5.58** | **1.95 - 15.92** | **0.001** |
| Young (2 - 4 years old) | 30 | 97 | 30.93% | 22.6% - 40.7% | **2.87** | **1.02 - 8.08** | **0.046** |
| Old (5 - 7 years old) | 5 | 37 | 13.51% | 5.9% - 28% |  |  |  |
| Very old (>7 years old) | 21 | 44 | 47.73% | 33.8% - 62.1% | **5.84** | **1.92 - 17.78** | **0.002** |
| Age |  |  |  |  |  |  |  |
| Age |  | 243 |  |  |  |  | <0.001 |
| Age<sup>2</sup> |  | 243 |  |  |  |  | <0.001 |
| Body Condition<sup>a</sup> |  |  |  |  |  |  |  |
| Body condition score |  | 182 |  |  |  |  | 0.0416 |
| Sett group size |  |  |  |  |  |  |  |
| Total |  | 251 |  |  |  |  | 0.814 |
| Adult |  | 251 |  |  |  |  | 0.434 |
| Cub |  | 251 |  |  |  |  | 0.086 |
| Cub percentage per sett |  |  |  |  |  |  |  |
| Low (<30%) | 58 | 191 | 30.37% | 24.3% - 37.2% |  |  |  |
| High (>30%) | 32 | 60 | 53.33% | 40.9% - 65.4% | **2.62** | **1.23 - 4.83** | **0.002** |
| Lactational status<sup>cd</sup> |  |  |  |  |  |  |  |
| Not Lactated | 3 | 10 | 30.00% | 10.8% - 60.3% |  |  |  |
| Lactated | 6 | 18 | 33.33% | 16.3% - 56.3% | 1.17 | 0.22 - 6.21 | 1 |

a: Only adults were included in this analysis b: Only females captured in spring were included in this analysis c: Fisher exact test

**Figure 1:**
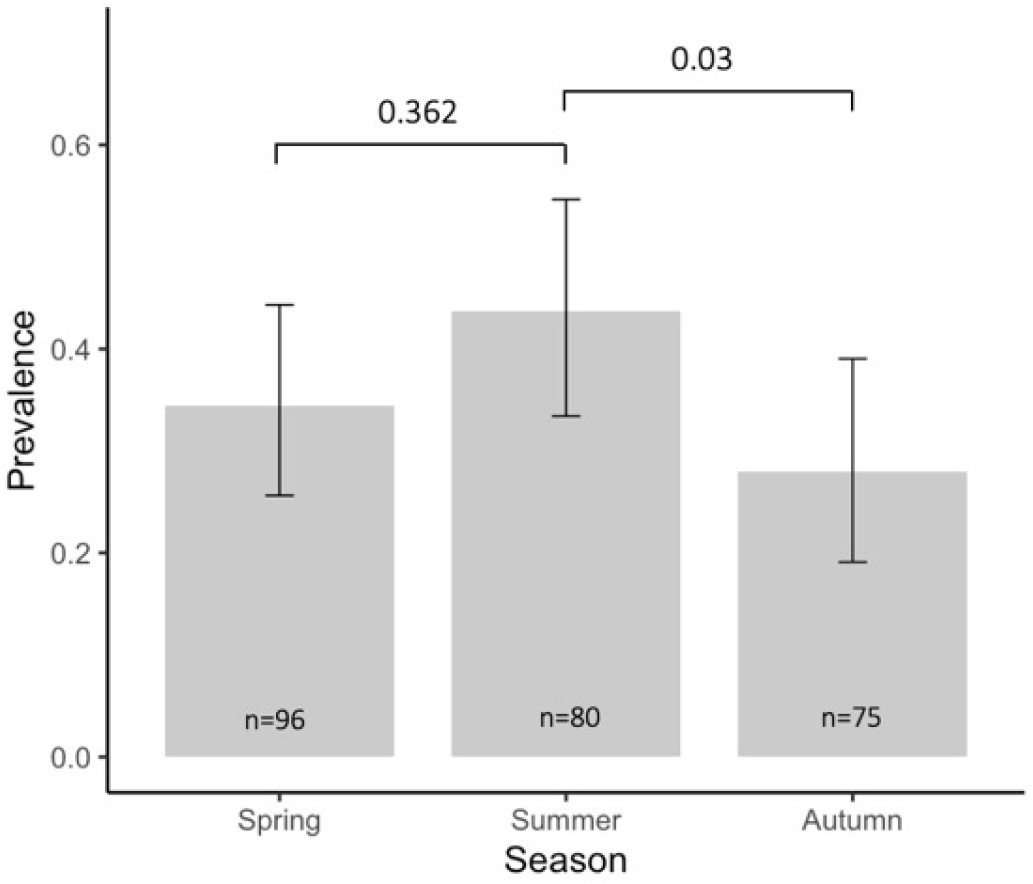
Difference of genital MusGHV-1 prevalence between seasons (Logistic regression analysis)

### Age effects on seasonal and sex-specific patterns of genital MusGHV-1 reactivation

There was strong evidence for an effect of age on genital MusGHV-1 where prevalence followed a U-shaped age curve, being lowest for badgers at the age of 5 or 6 years old (Figure 2, quadratic term, adjusted R^2^= 0.683, F-statistic= 12.83 on 2 and 9 DF, p-value= 0.002). When divided by age classes, prevalences in juveniles (46.6%, 34/73) and very old badgers (47.7%, 21/44) were higher than young (30.9%, 30/97) and old (13.5%, 5/37) adults (Table 2). However, no effect of sex (logistic regression analysis, p=0.679) was observed in the univariate analysis.

**Figure 2:**
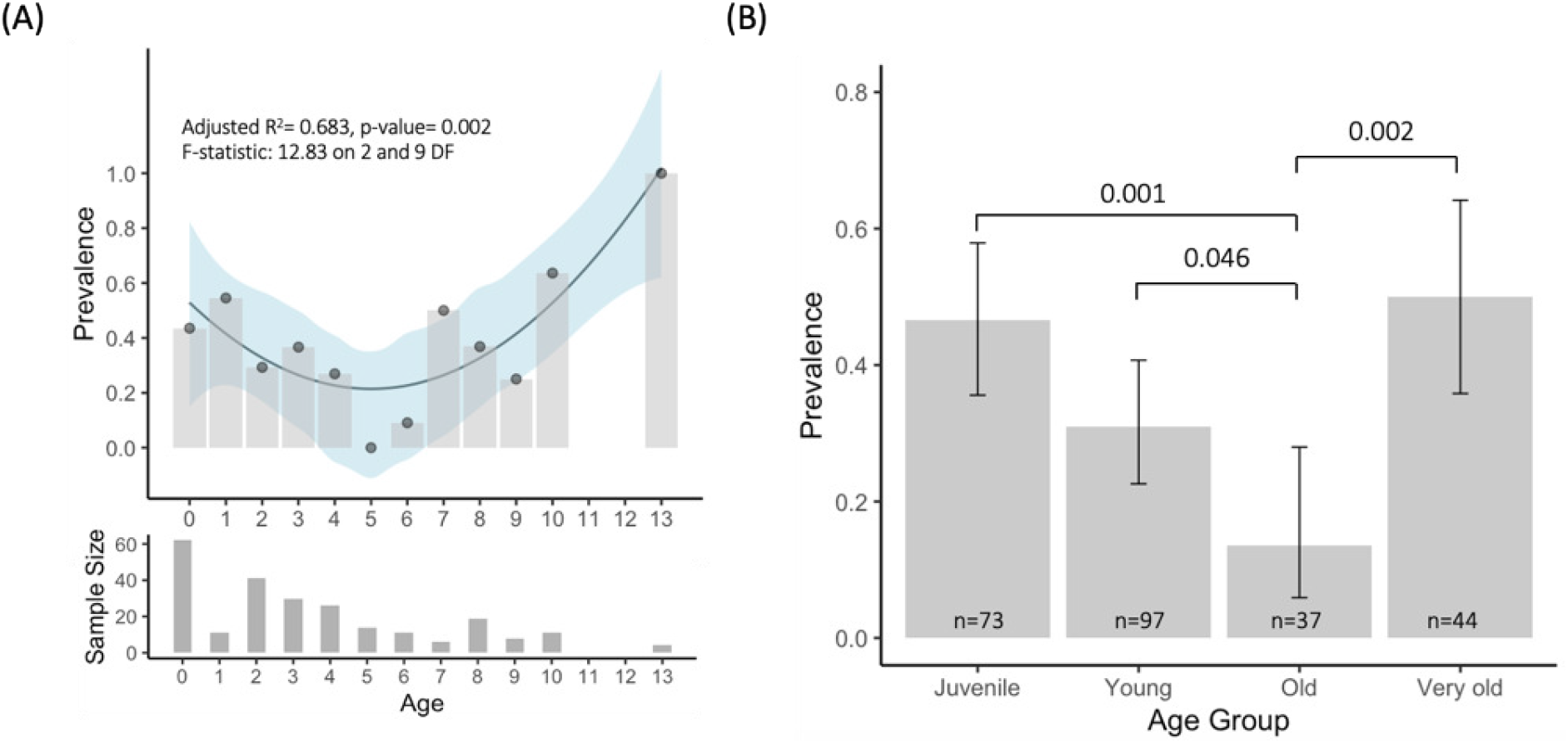
Difference of genital MusGHV-1 prevalence between exact age (A) and age groups (B) (Logistic regression analysis)

### BCS effects on seasonal and sex-specific patterns of genital MusGHV-1 reactivation

Although adults with lower BCS had a higher probability of genital MusGHV-1 reactivation according to our univariate analysis (p-value=0.022, Table 2), when grouped by seasons and sex this relationship was only significant in adult males in summer (Kruskal-Wallis tests, p-value = 0.006) (Figure 3).

**Figure 3:**
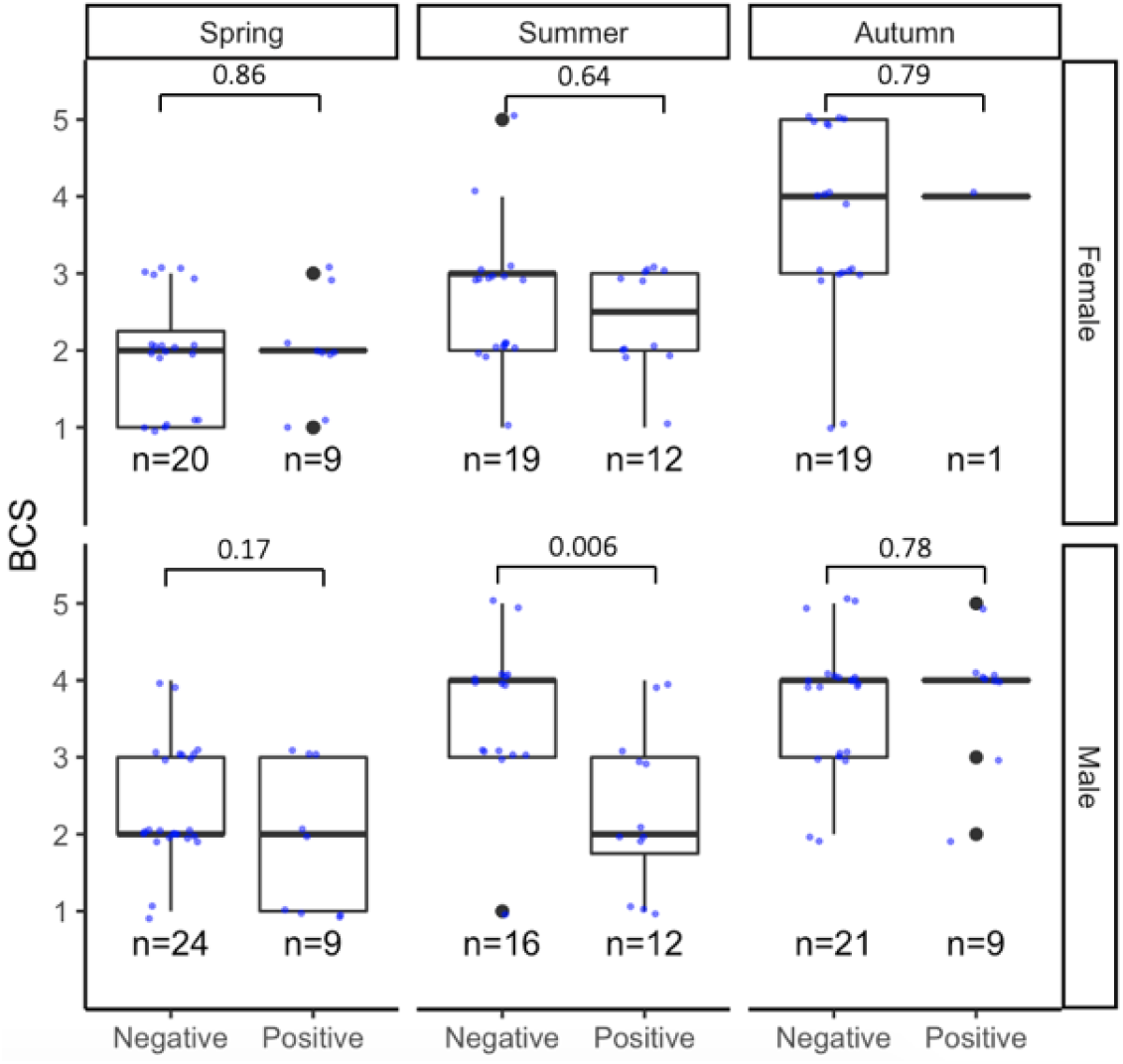
Comparison of body condition score difference of adults between genital MusGHV-1 reactivation status. The p-values of each Kruskal-Wallis test results are showing above each box plot.

### Group effects on seasonal and sex-specific patterns of genital MusGHV-1 reactivation

Genital MusGHV-1 reactivation was significantly more prevalent in females living in setts comprised by >30% cubs (logistic regression analysis, p-value = 0.002), but there was no evidence for the absolute number of adults, cubs nor the total number of badgers resident in each sett affecting reactivation rates (Table 2).

### Effects of recent reproductive success on female genital MusGHV-1 prevalence

There was no evidence for recent lactation affecting the reactivation rate in sexually mature females in spring (Fisher’s exact test, p-value=1) (Table 2).

### Multivariable analysis of genital MusGHV-1 reactivation

Because the univariate analysis indicated that juveniles and adults appear to have different patterns of reactivation, we separated these data to generate multivariable models of juveniles (n=73) and adults (n=178). We included all variables initially for which there was any evidence for an effect in the univariate analyses (p-value <0.10, Table 2). The most parsimonious models for juveniles (Table 3) and adults (Table 4) both had diagnostically acceptable (53) AUC area of 0.748 and 0.726 (Figure S3 and S4), respectively. In juveniles, multivariable analysis showed that all females, and all juveniles with higher BCS, were at particular risk of genital MusGHV-1 reactivation (Table 3). In adults, in contrast, males are at a higher risk of MusGHV-1 reactivation than are females, where all adults experience the highest reactivation risk in summer, and older (≥8 years) badgers, and female living in setts with a higher percentage (over 30%) of cubs are at particular risk (Table 4).

**Table 3:** Final general mixed effect model of multivariable logistic regression analysis for juveniles Formula: MusGHV ~ Sex + Body condition + (1|Tattoo); number of observations: 72; groups by tattoo number: 48

| Group | Estimate | Standard error | z value | Adjusted OR | 95% CI | p value |
| --- | --- | --- | --- | --- | --- | --- |
| (Intercept) | -1.226 | 0.7851 | -1.562 |  |  | 0.1184 |
| Sex |  |  |  |  |  |  |
| Female |  |  |  |  |  |  |
| Male | -1.0519 | 0.5123 | -2.054 | **0.35** | **0.13 - 0.95** | **0.04** |
| Body Condition |  |  |  |  |  |  |
| BCS | 0.6245 | 0.2876 | 2.172 | **1.87** | **1.06 - 3.28** | **0.0299** |

**Table 4:** Final general mixed effect model of multivariable logistic regression analysis for adults Formula: MusGHV ~ Sex + Season + AgeGroup + Cub percentage + Sex*Cub percentage + (1|Tattoo); Number of observations: 178; Groups by tattoo number: 101

| Group | Estimate | Standard error | z value | Adjusted OR | 95% CI | p value |
| --- | --- | --- | --- | --- | --- | --- |
| (Intercept) | -0.9952 | 0.4859 | -2.048 |  |  | 0.04 |
| Sex |  |  |  |  |  |  |
| Female |  |  |  |  |  |  |
| Male | 0.8171 | 0.4126 | 1.98 | **2.26** | **1 - 5.1** | **0.048** |
| Season |  |  |  |  |  |  |
| Spring | -1.037 | 0.4255 | -2.437 | 1.61 | 0.66 - 3.91 | 0.297 |
| Summer | -2.2692 | 0.6333 | -3.583 | **3.08** | **1.27 - 7.47** | **0.013** |
| Autumn |  |  |  |  |  |  |
| Age class |  |  |  |  |  |  |
| Young | -1.1084 | 0.468 | -2.368 | **0.35** | **0.15 - 0.82** | **0.015** |
| Old | -2.4001 | 0.7052 | -3.404 | **0.1** | **0.03 - 0.36** | **<0.001** |
| Very old |  |  |  |  |  |  |
| Cub percentage per sett |  |  |  |  |  |  |
| Low (<30%) |  |  |  |  |  |  |
| High (>30%) | 1.8903 | 0.6968 | 2.713 | **6.62** | **1.69 - 25.95** | **0.006** |
| Interaction |  |  |  |  |  |  |
| Female:Cub percentage |  |  |  |  |  |  |
| Male:Cub percentage | -2.3686 | 1.0492 | -2.258 | **0.09** | **0.01 - 0.73** | **0.024** |

### Genetic diversity of MusGHV-1 in the Wytham badger population

We sequenced 5 MusGHV-1 positive PCR products of a partial DNA polymerase gene. All sequences were trimmed to 694 base pairs and confirmed to be MusGHV-1 according to the NCBI online blasting service, returning 98.7% (n=3) and 100% (n=2) nucleotide identity to the published MusGHV-1 sequence isolated from a badger in Cornwall, England (Accession number: AF275657).

## Discussion

Herpesvirus reactivation triggered by stress has been widely confirmed naturally and experimentally by corticosteroid injection in humans and domestic animals (5, 14). Linking stress and viral reactivation in wildlife, however, is particularly challenging due to the difficulties of monitoring individual stress levels in the field, and typically this relationship can only be confirmed experimentally by taking subjects into captivity at least temporarily (15). Using indicators that have been linked to stress hormone levels in previous studies can thus provide an informative way to study the relationship between stress and herpesvirus reactivation in free-ranging wildlife.

Faecal corticosteroid measurements from badgers in Ireland (30) evidence higher stress levels in summer likely associated with dry environmental conditions that result in lower earthworm availability (i.e., the badgers’ main food type (46)); similarly, in our own study population, summer drought is an established mortality factor due to starvation/ malnutrition (32, 33). This corresponds with our finding that, in all adults, seasonal MusGHV-1 reactivation rates were highest in summer, but females tended to have higher viral reactivation levels than males in spring – possibly due to reproductive stresses, while the reverse was true in autumn (Figure 1) (30). Interestingly, however, we found no correlation between BCS and MusGHV-1 reactivation across all seasons in females, implying that reduced body condition was not necessarily indicative of physiological stress. In fact, Bright Ross et al. (subm) found complex relationships between body-condition, survival and reproductive success in this same population, where although breeding females lose condition, they often end up being no thinner than non-breeding females as they were in much better condition in winter (i.e. before pregnancy/ lactation).

From the perspective of male rates of reactivation, males with higher testosterone levels tend to be thinner during spring and summer (54), but tend to mate more often (55). In our recent survey of Irish badger populations (20), the high rate (over 80%) of genital MusGHV-1 reactivation in adult males during the peak (postpartum) mating season, from mid-January to mid-February, implies not only a link to mating activity, but also a mechanism enhancing transmission. This corroborates another finding in the same study that males with more spermatozoa have a higher prevalence of genital MusGHV-1 reactivation (20); linking higher sexual activity to higher STI prevalence as reported also in many other studies (56–59). We were unable to explicitly test effects relating to badger mating behaviour in our study because we can not trap during late pregnancy and neonatal cub care, to avoid stressing mothers or depriving cubs of maternal care. Nevertheless, although reactivation rate in autumn, when food sources are most abundant, and badgers undergo a period of reproductive quiescence (60), and thus experience less implicit stress, was significantly lower compared only to summer, but not to spring. This suggests that other factor(s) (e.g. sex hormone cycles (61), oxidative stress (62), genital microbiome (63) or bacterial co-infections) might also be affecting reactivation rates, beyond the scope of our current study.

In terms of age class effects, the high genital reactivation rate detected in cubs and yearlings suggests that badgers contract MusGHV-1 early in life, before reaching sexual maturity. Although MusGHV-1 reactivates repeatedly throughout life, reactivation tends to be less frequent in young and old adults compared to adults in their prime, although rates increase in very old individuals. This matches patterns in humans where most people become infected with herpes during their childhood/ adolescence (e.g. 100% and 70% seroprevalence of EBV before age 14 in Hong Kong and the United Kingdom (64)), then typically experience viral latency during their prime, but can suffer from increasingly longer and more frequent herpesvirus reactivation that sometimes cause mild disease (e.g., shingles (65) in old age due to lowered immune response (66, 67)).

Since vertical transmission of MusGHV-1 through the placenta is unlikely (20), and the potential for infection from the vaginal tract during parturition is equally low due to low genital MusGHV-1 reactivation rate in pregnant females (20), we hypothesise that cubs contract primary infection through close contact with virus-shedding conspecifics (20). Thereafter genital reactivation in cubs may arise after primary acute infection through non-sexual routes and subsequent latency, as observed in the murine model where *Murine herpesvirus 4* (MuHV-4, also a gammaherpesvirus), inoculation in the nasal cavity results in acute infection in the respiratory tract and lungs and establishes latency in the spleen, but then reactivates in the vaginal tract 17-21 days after inoculation (61). MuHV-4 nasal cavity inoculation, however, does not result in reactivation in male genital tracts, and transmission is only possible from females to males. After sexual intercourse with virus-shedding female mice, the virus then replicates in the male penis for 3 weeks. Interestingly, also in badgers, female juveniles are at higher risk of MusGHV-1 reactivation in the genital tract than are males. Furthermore, juveniles in better body condition, regardless of sex, exhibit higher reactivation prevalence. Indeed, juvenile males in better body condition enter puberty earlier than thinner males (11 months compared to 22 – 28 months: (57)). Once juveniles enter puberty they will experience an increased risk of contracting MusGHV-1 through sexual contact and/ or that their latent infection is reactivated through mating resulting in viral shedding in the genital tract.

Our results also show that social group structure can affect prevalence of genital herpesvirus reactivation, particularly the proportion of cubs within a residential group. This trend was more apparent in females than in males. This may be because badger cubs generally carry higher pathogen burdens than adults (68, 69), and thus increase per capita immunity burden among all badgers resident in the respective sett (64).

## Conclusion

Our study demonstrates - for the first time in the wild - the link between stress experienced by the host and latent virus reactivation. Amplified stress levels induced by human disturbance as well as food insecurity and more frequent catastrophic weather events arising from human induced rapid environmental change (HIREC) could therefore not only increase the risk of disease development, promotion of transmission within a population, but also negatively impact host reproductive fitness through latent virus reactivation in the reproductive tract (19). Careful monitoring of endemic latent virus infection as well as surveillance for possible newly emerging strains should be included when planning *in situ* and *ex situ* conservation programmes for endangered species (70).

## Acknowledgments

The authors would like to thank Dr. Nadine Sugianto, Dr. Sil van Lieshout, Dr. Tanesha Allen, and Julius Bright-Ross for assistance with sample collection, fieldwork and MNA data provision. MST would like to thank the Ministry of Education in Taiwan and Lady Margaret Hall, University of Oxford, for scholarship support. The authors also thank Paul Johnson and Ta-Chun Liu for providing advice on statistics. The authors claim no conflict of interests in the present work.

## Author Contributions

Project conception: CBD,MST; samples collection: CBD,CN,MST; laboratory work: MST; data analysis: MST; writing and revision: MST,CBD,SF,DWM; All authors have read and approved the manuscript.

---

**Figure S1:**
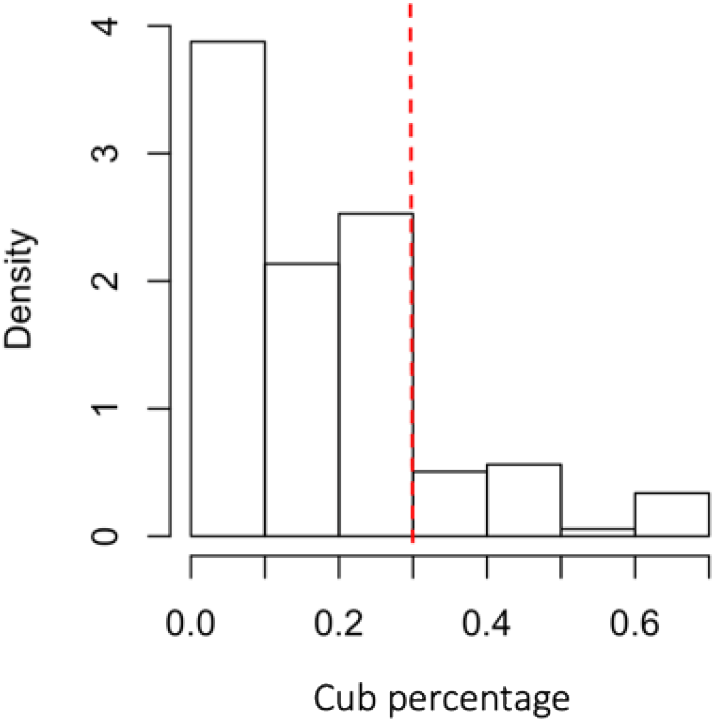
Data distribution of cub percentage (n=251). The cutoff point 30% (red dashed line) is used to divide the tail and the distribution in the left.

**Figure S2:**
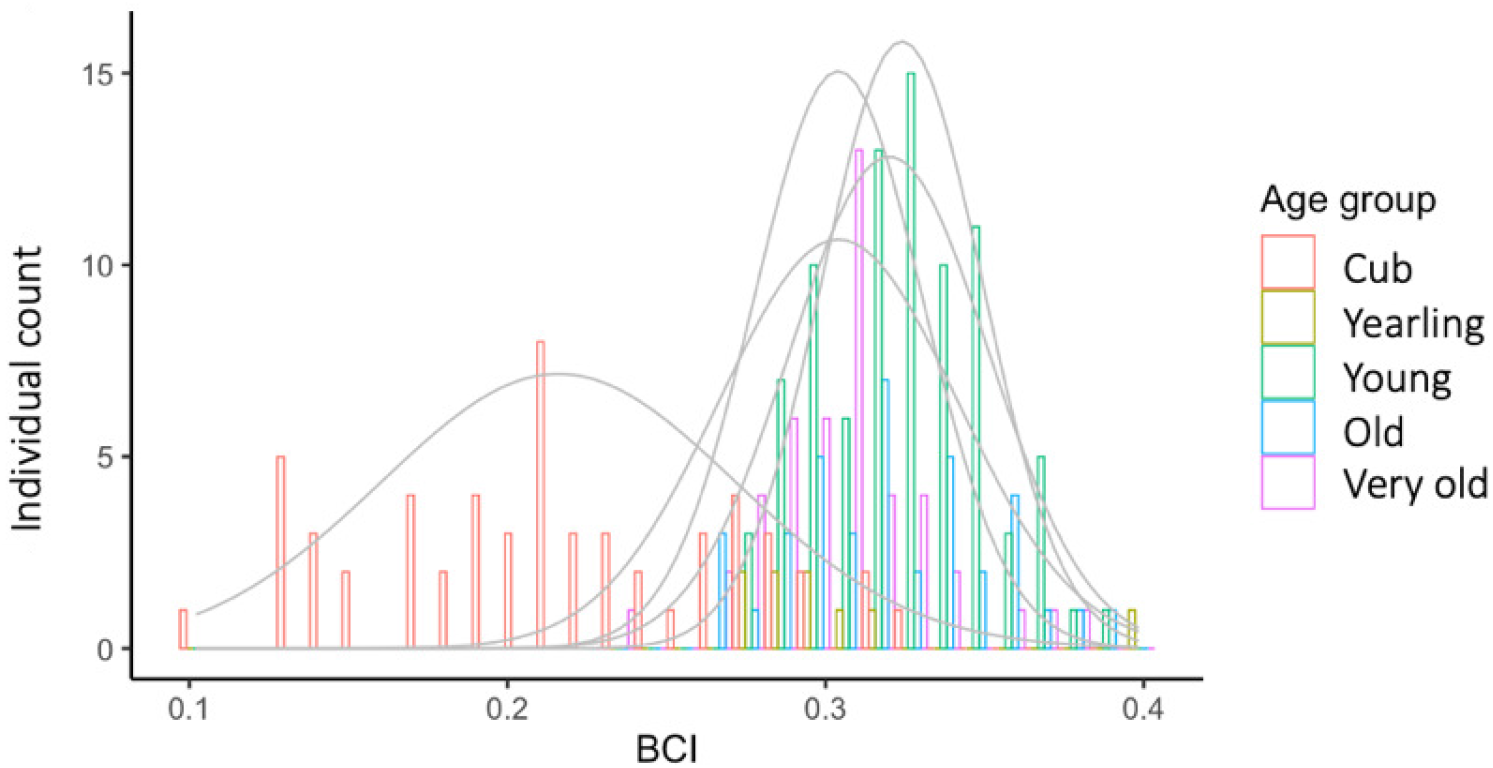
BCI data distribution by age groups showing distinct distribution of cub BCI comparing with other groups (A)

**Figure S3:**
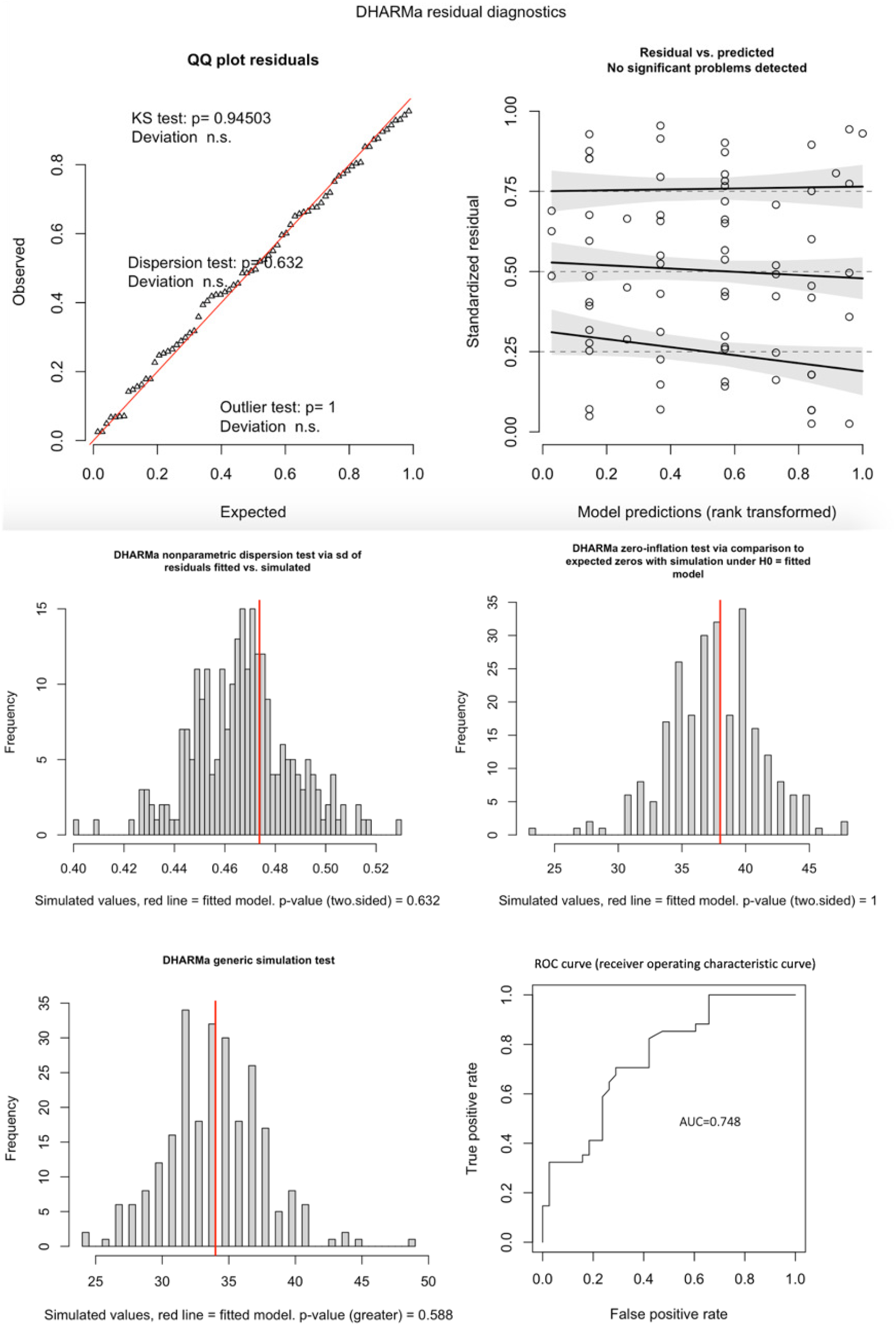
Diagnostic plot of the final multivariable logistic regression model of juvenile genital MusGHV-1 reactivation

**Figure S4:**
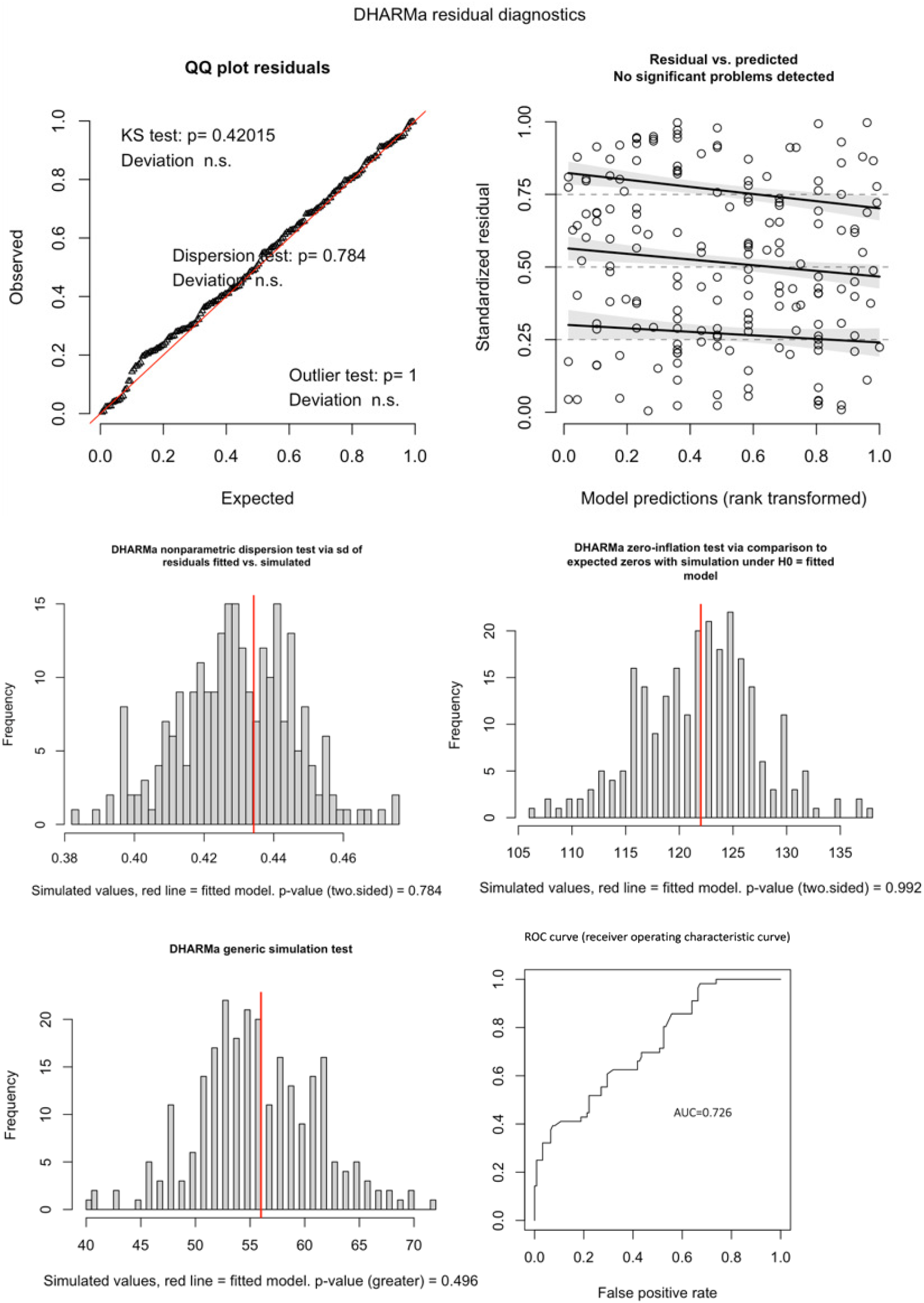
Diagnostic plot of the final multivariable logistic regression model of adult genital MusGHV-1 reactivation

